# Oleic acid triggers CD4^+^ T cells to be metabolically rewired and poised to differentiate into proinflammatory T cell subsets upon activation

**DOI:** 10.1101/2024.02.16.580665

**Authors:** Nathalie A. Reilly, Friederike Sonnet, Koen F. Dekkers, Joanneke C. Kwekkeboom, Lucy Sinke, Stan Hilt, Hayat M. Suleiman, Marten A. Hoeksema, Hailiang Mei, Erik W. van Zwet, Bart Everts, Andreea Ioan-Facsinay, J. Wouter Jukema, Bastiaan T. Heijmans

## Abstract

T cells are the most common immune cells in atherosclerotic plaques and the function of T cells can be altered by fatty acids. Here, we show that pre-exposure of CD4^+^ T cells to oleic acid, an abundant fatty acid linked to cardiovascular events, results in a preferential differentiation into pro-inflammatory subsets upon activation by upregulating core metabolic pathways. RNA-sequencing of non-activated CD4^+^ T cells revealed that oleic acid upregulates genes encoding enzymes responsible for cholesterol and fatty acid biosynthesis. Transcription footprint analysis linked this rewiring to the differentiation of pro-inflammatory subsets. Indeed, spectral flow cytometry showed that pre-exposure to oleic acid results in a skew toward IL-9, IL-17A, IL-5 and IL-13 producing T cells upon activation. Importantly, inhibition of either cholesterol or fatty acid biosynthesis abolishes this effect, suggesting a beneficial role for statins beyond cholesterol lowering. Taken together, fatty acids may affect inflammatory diseases by influencing T cell metabolism.

## Introduction

Atherosclerosis is the primary underlying cause of cardiovascular disease and is driven by the interactions between the immune system, lipids, and the vascular wall^1, 2^. Recent single-cell RNA sequencing and mass cytometry studies showed that T cells make up the majority of immune cells in atherosclerotic plaques, half of which are CD4^+^ T cells^3–6^. This indicates that the role of CD4^+^ T cells in atherosclerosis is much greater than previously recognized^7–9^. Lipids and in particular fatty acids are known to have a major influence on the function of CD4^+^ T cells^2^. While previous research evaluated the effect of fatty acids on CD4^+^ T cell function during or after activation^2, 10–13^, interactions between fatty acids and CD4^+^ T cells relevant for atherosclerosis can already occur when the cells are in a non-activated state^2^. While the impact of these interactions have not been studied, they may skew the differentiation towards pro- or anti-inflammatory subsets^8, 14^ once the CD4^+^ T cells infiltrate atherosclerotic plaques or other disease sites such as rheumatoid arthritis, and become activated.^2, 15^

Fatty acids affect CD4^+^ T cells in multiple ways ranging from activation, proliferation, to differentiation^2^. It is thought that these effects are largely mediated by changes in metabolism^16^. In a non-activated state, like in the circulation, CD4^+^ T cells rely on oxidative phosphorylation and β-oxidation of fatty acids for energy production^17, 18^. However, upon activation, CD4^+^ T cells switch their metabolism to fatty acid biosynthesis and aerobic glycolysis to support cell growth and proliferation, reminiscent of the Warburg effect^18–20^. Importantly, the generation of specific T cell subset populations is associated with this metabolic reprogramming ^10, 21–26^. The generally pro-inflammatory^14, 27, 28^ T helper 1 (T_H_1), and T helper 17 (T_H_17) cells, but also T helper 2 (T_H_2) cells that can be both pro- and anti-inflammatory, rely on pathways of aerobic glycolysis upon activation^29–35^. In contrast, the generally anti-inflammatory regulatory T (T_reg_) cells mainly remain reliant on oxidative phosphorylation even after activation, indicating that the metabolic state of the cell may influence T cell effects in disease^18, 22, 23, 34^. Therefore, fatty acid-mediated metabolic reprogramming of CD4^+^ T cells may affect the initiation and progression of atherosclerosis by skewing CD4^+^ T cells towards a pro-inflammatory phenotype.

In this study, we characterized the effects of oleic acid on non-activated CD4^+^ T cell function. Oleic acid is a monounsaturated fatty acid that is of particular interest since it is one of the most abundant fatty acids in the circulation^36^ and is independently associated with an increased risk of cardiovascular events^37^. To do so, we performed RNA sequencing and an 850k DNA methylation array on non-activated CD4^+^ T cells exposed to oleic acid at 5 different time points. Furthermore, we performed spectral cytometry post-activation for various CD4^+^ T cell markers. We find that oleic acid exposure leads to a metabolic reprogramming and generates a profile that becomes skewed towards T_H_2, T_H_17 and, notably, T_H_9 CD4^+^ T cells after activation. This skewed profile post-activation is blocked by the addition of metabolic inhibitors during the initial oleic acid exposure.

## Results

### Establishing a model to study the effect of oleic acid on non-activated CD4^+^ T cells

Prior to studying the effect of oleic acid on non-activated CD4^+^ T cells, we evaluated various experimental conditions in order to establish an *in vitro* exposure model. The cellular response to oleic acid was assessed by measuring cell viability and the expression of *CPT1A*. The *CPT1A* gene encodes the long chain fatty acid transporter carnitine palmitoyl transferase 1a, a rate limiting enzyme in the metabolic process of β-fatty acid oxidation. First, three different types of culturing conditions for non-activated CD4^+^ T cells were compared: 5% FCS medium with oleic acid bound to fatty acid free (FAF) bovine serum albumin (BSA), 5% FCS medium with oleic acid diluted in 5% FCS medium, and FAF medium with oleic acid bound to FAF BSA. However, the latter two conditions led to either undissolved oleic acid or a low cell viability (Supp. Fig. 1a). The first condition produced the largest *CPT1A* response while maintaining a high cell viability (Supp. Fig. 1), presumably because oleic acid bound to BSA and the presence of FCS may be a better approximation of physiological conditions. In addition, various oleic acid concentrations used in previous studies were evaluated^10–12, 38–42^. A concentration of 30μg/mL was observed to result in the highest *CPT1A* upregulation (9.83 fold, SE 5.60) while maintaining cell viability (84.36%, SE 0.49%; Supp. Fig. 1). Importantly, this concentration does not exceed the concentration of oleic acid in the human circulation (0.03 to 3.2 mmol/L or 84.8 to 903.9 μg/mL)^43^. The oleic acid solvent ethanol did not influence the results (Supp. Fig. 1). Finally, we measured the oleic acid concentration in 5% FCS using the SLA method. On the basis of these measurements, we calculated that the culture medium with 5% FCS contained 0.26 μg/mL of free oleic acid and 4.39 μg/mL oleic acid as components of larger molecules including cholesterol esters and sphingolipids.

### Transcriptomic analysis of oleic acid exposed non-activated CD4^+^ T cells

In order to identify the molecular features that define the effect of oleic acid exposure on non-activated CD4^+^ T cells *in vitro*, we exposed non-activated CD4^+^ T cells to 30μg/mL oleic acid for 0.5, 3, 24, 48, or 72h (n=9; Fig. 1a). First, we measured *CPT1A* expression and found that its expression consistently increased over time indicating a robust response to oleic acid exposure across donors, while *CPT1A* expression did not change under control conditions (Fig. 1b). Next, we analyzed the transcriptome of non-activated CD4^+^ T cells after oleic acid exposure using RNA-seq. Oleic acid induced differential expression of 544 genes (P_FDR_ < 0.05) that clustered into 310 upregulated genes and 234 downregulated genes (Fig. 1c; Supp. Fig. 2; Supp. Table 1a and b). There was no statistical evidence for further subdivisions of the two clusters, for example in fast and slow-responding genes.

**FIGURE 1.**
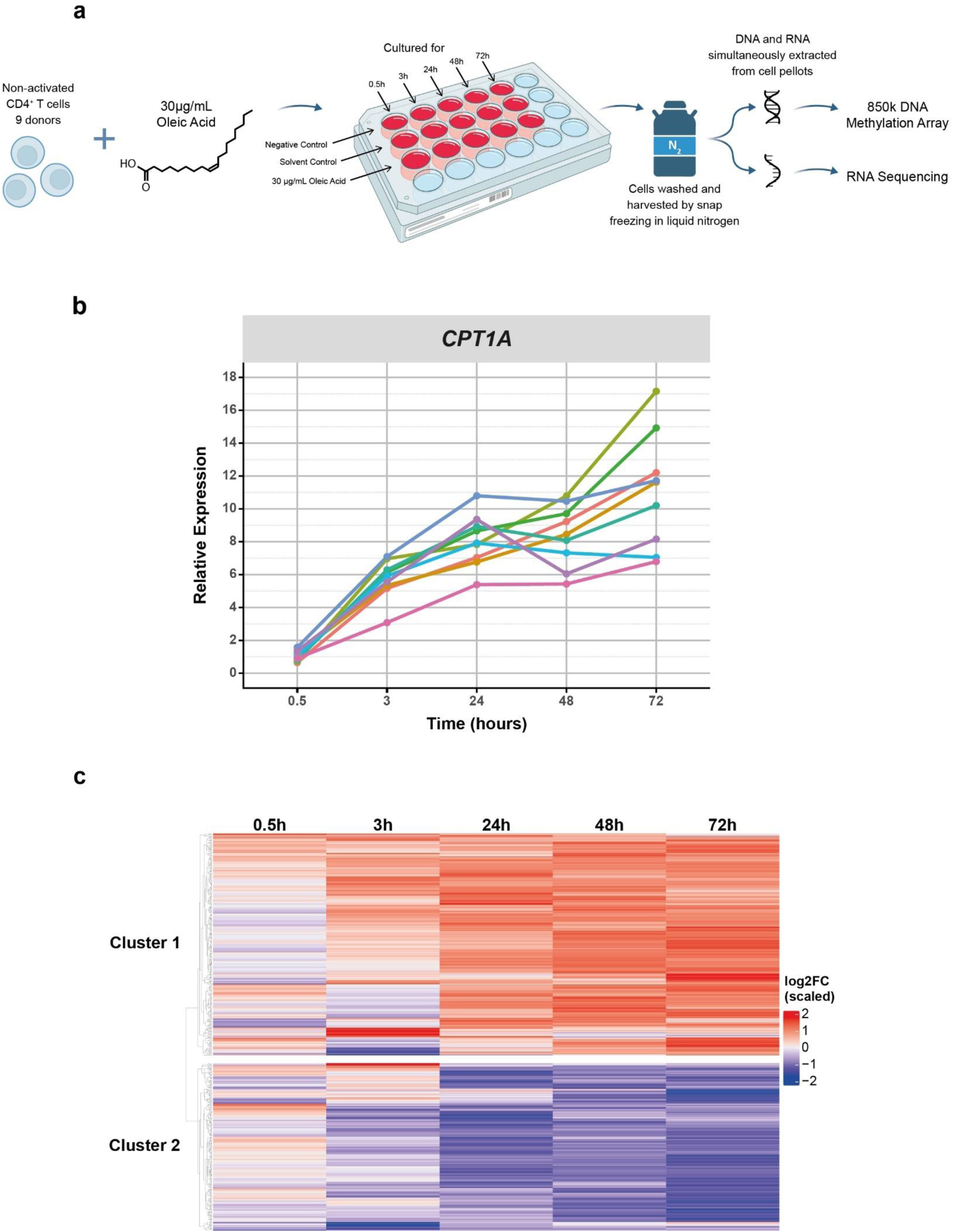
**(a)** Experimental set up for RNA sequencing and DNA methylation measurements of oleic acid exposed non-activated CD4^+^ T cells, n =9. **(b)** Line plot showing the relative expression of *CPT1A* per donor across time as a confirmation of the *in vitro* model by RT-qPCR. Values are colored by donor across time. On average *CPT1A* was upregulated 1.03 SE 0.10 fold at 0.5h, 5.73 SE 0.40 fold at 3h, 8.08 SE 0.53 fold at 24h, 8.39 SE 0.62 fold at 48h, and 11.09 SE 1.16 fold at 72h, n =9. **(c)** Differentially expressed genes (DEGs) in oleic acid exposed non-activated CD4^+^ T cells across time. Heatmap obtained from the DESeq2 analysis resulting in 544 DEGs (PFDR < 0.05). DEGs were plotted across time to show the genes expression as log2FoldChange at each time point. Unsupervised K-means clustering indicated 2 clusters. Cluster 1 contains 310 of the DEGs, which are generally upregulated and are represented in red and cluster 2 contains 234 of the DEGs, which are generally downregulated and are represented in blue, n = 9.

We first examined the functions of the 310 genes that were upregulated in non-activated CD4^+^ T cells by oleic acid exposure. We inspected the top differentially expressed genes (Supp. Fig. 2a). The top-differentially expressed gene was *CPT1A* highlighting the involvement of β-fatty acid oxidation. In addition, we found an increased expression of *HMGCR* (3-hydroxy-3-methyl-glutaryl-coenzyme A reductase), encoding the rate-limiting enzyme for cholesterol biosynthesis and *ACACA* (acetyl-coenzyme A carboxylase 1), encoding the rate-limiting enzyme of fatty acid biosynthesis. Furthermore, transcripts of several aerobic glycolysis related genes, such as *TKT* and *PGD*, were upregulated (Supp. Fig. 2a). A formal analysis of enriched biological processes among all 310 upregulated genes confirmed the involvement of metabolism. In particular, cholesterol biosynthesis (P_FDR_ < 0.001), homeostasis (P_FDR_ < 0.001), and signaling of mTORC1 (P_FDR_ < 0.001), a key complex of mTOR which aids in the switch towards aerobic glycolysis and fatty acid biosynthesis, were enriched (Fig. 2a). Mapping the upregulated genes to canonical metabolic pathways further supported a specific metabolic rewiring of oleic acid exposed non-activated CD4^+^ T cells (Fig. 2b). First, oleic acid can first be catabolized through beta oxidation to produce acetyl CoA, which can then be used as a starting point for cholesterol and fatty acid biosynthesis. In addition to *CPT1A*, we found 4 out of 15 enzymes in β-fatty acid oxidation (including *SLC25A20*, *ACADVL*, *ACAA2*) and 2 out of 15 enzymes in the aerobic glycolysis pathway to be upregulated (*TKT* and *PGD*; Fig. 2b; Supp. Fig. 3 and 4). Remarkably, on top of *HMGCR*, 15 out of 20 enzymes involved in cholesterol biosynthesis were upregulated in our gene set, including several key rate limiting genes (such as *HMGCS1*, *SQLE*, *MVD*, and *MVK*). More specifically, 9/11 components of the mevalonate, 6/9 of the Bloch and, 6/9 of the Kandutsch-Russell pathway, together responsible for cholesterol biosynthesis, were upregulated (Fig. 2b; Supp. Fig. 5). The upregulated gene set also included *ACACA* and *FASN* that encode the two enzymes that together are responsible for the 37 reactions making up fatty acid biosynthesis (Fig. 2b; Supp. Fig. 6). Of note, the genes *ACACA* and *FASN* have been implicated in the differentiation towards T_H_17 cells, a highly pro-inflammatory subset of CD4^+^ T cells^42^. Furthermore, aerobic glycolysis, cholesterol and fatty acid biosynthesis are the hallmark metabolic processes of activated T cells and suggest that non-activated CD4^+^ T cells undergo a metabolic reprogramming upon oleic acid exposure that may poise the cells for a different response to activation.

**FIGURE 2.**
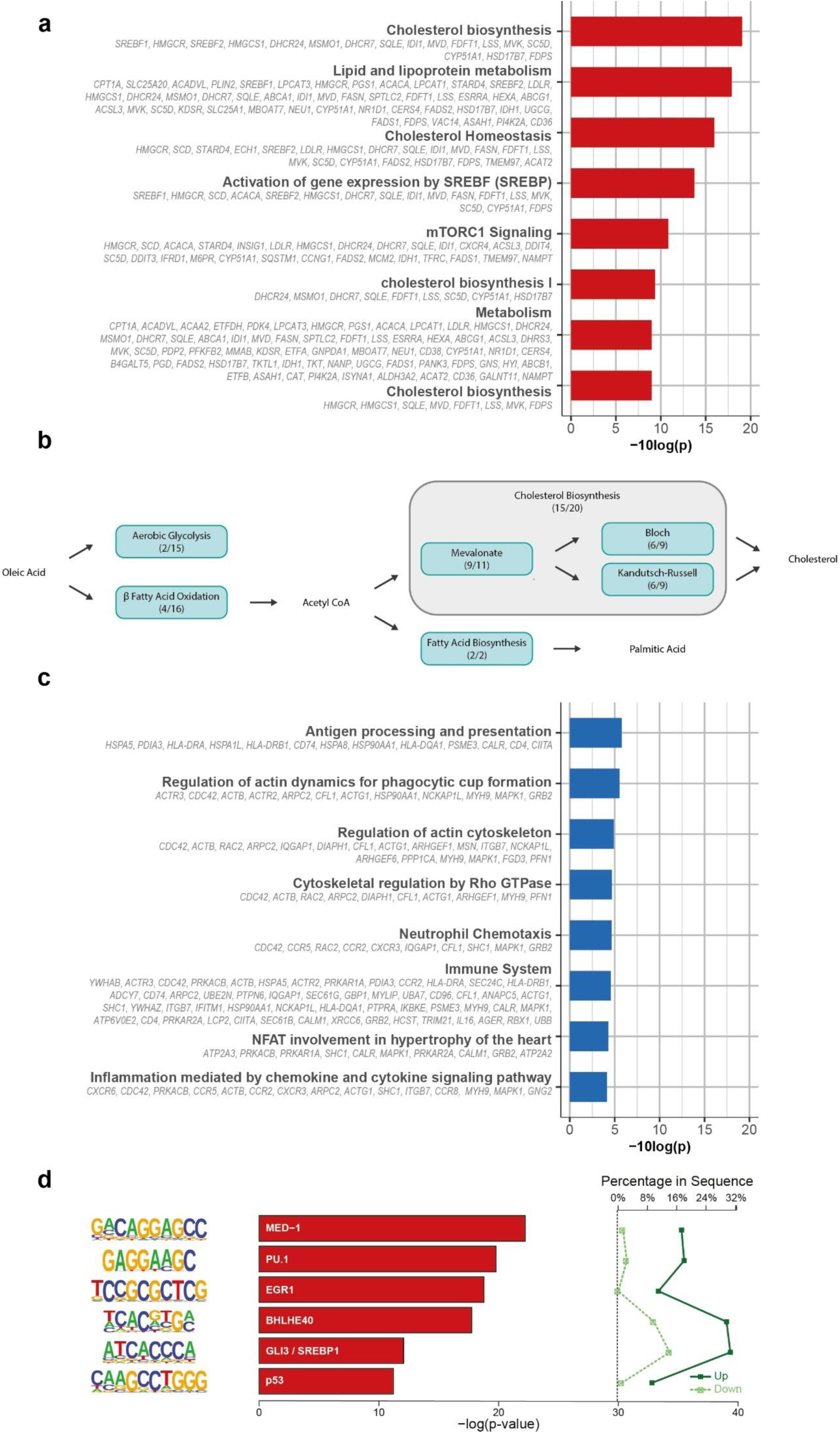
**(a)** Pathway enrichment analysis of cluster 1 DEGs generated using *clusterProfiler* using 10 human pathway databases. Top 8 enrichments are shown. **(b)** Illustration of canonical pathway map of oleic acid metabolism by non-activated CD4^+^ T cells exposed to oleic acid. Blue boxes indicate metabolic pathways with the number of genes present in that particular pathway from cluster 1 of the RNA sequencing. Cholesterol biosynthesis can be divided into 3 separate pathways indicated by the surrounding grey rectangle. **(c)** Pathway enrichment analysis of cluster 2 DEGs generated using *clusterProfiler* using 10 human pathway databases. Top 8 enrichments are shown. **(d)** *de novo* motif analysis on promotors of up versus down regulated genes. Enrichment of transcription factor binding motifs was performed using HOMER. 6 motifs are shown with supplementing information on p-value, percentage of genes in upregulated gene set and percentage of genes in downregulated gene set, transcription factor name, -log(p-value), and percentage in sequence.

We then examined the functions of the 234 genes that were downregulated in non-activated CD4^+^ T cells by oleic acid exposure. We first inspected the top differentially expressed genes (Supp. Fig. 2b). Among the top downregulated genes, decreased expression of *CXCR6* and *CCR5*, important chemokine receptors in the T cell immune response, were measured. Moreover, expression of *TPM4*, encoding actin-binding proteins involved in the cytoskeleton, and *DMTN*, encoding an actin binding and bundling protein that stabilizes the actin cytoskeleton, were also downregulated. A formal analysis of the enriched biological processes among all 234 downregulated genes revealed a wide variety of different pathways. In line with the genes observed among the top-downregulated genes, this included processes involved in immune response (*CCR2*, *CCR8, HLA-DRA*, *SLC2A1*) (P_FDR_ < 0.001) and actin cytoskeleton organization (*ACTB*, *RAC2*, *ARPC2*, *IQGAP1*) (P_FDR_ < 0.001) (Fig. 2c). In addition, processes involved in chemotaxis (P_FDR_ < 0.001), chemokine and cytokine signaling (P_FDR_ < 0.001), and Rho GTPase regulation (P_FDR_ < 0.001) were also downregulated (Fig. 2c). Overall, these data point to a broad yet a-specific downregulation of genes in oleic acid exposed non-activated CD4^+^ T cells, perhaps to cope with the influx of the fatty acid.

Next, we investigated whether specific transcription factors may underlie the differential expression observed by testing the enrichment of transcription factor binding motifs in upregulated vs downregulated genes. The top motifs enriched among up-regulated genes included key transcription factors PU.1, EGR1, BHLHE40, and SREBP1 (Fig. 2d). Notably, PU.1 is the key transcription factor for the development of T_H_9 cells. BHLHE40 has been linked to T_H_17 development and pathogenicity in autoimmune encephalomyelitis suggesting an additional possible preference towards T_H_17 differentiation post-activation^44, 45^. Furthermore, EGR1 and SREBP1 are either involved in the activation of Tbet or fatty acid and cholesterol biosynthesis, respectively^46, 47^. These data further support the notion that oleic acid exposed non-activated CD4^+^ T cells may be poised to differentiate towards pro-inflammatory T cell subsets, including T_H_9 and T_H_17, after activation.

In addition to a transcriptomics analysis, genome-wide DNA methylation was analyzed to explore whether oleic acid exposure was indicative of a shift in the epigenetic landscape of the cells. However, no evidence for oleic acid-induced epigenetic changes at the level of DNA methylation were observed in non-activated CD4^+^ cells (P_FDR_ > 0.05, Supp. Table 1c and d). Further inspection showed that neither any of the top-10 differentially methylated CpGs mapped to genes involved in metabolism or T cell function, nor was any of the nearest genes among the genes found to be differentially expressed (Supp. Table 1e and f). These results indicate that changes in the transcriptome were not mirrored by similar changes in the methylome within 72h of exposure to oleic acid.

### Oleic acid induced CD4^+^ T cell phenotypes after activation

To determine the functional impact of the transcriptomic changes identified, we characterized the phenotypes of CD4^+^ T cell that were pre-exposed to oleic acid or control conditions and subsequently activated in the absence of oleic acid. To this end, non-activated CD4^+^ T cells of 8 out 9 donors for whom sufficient cells were available, were again exposed to 30μg/mL oleic acid (Fig. 3a). The effect of exposure was confirmed by an upregulation of *CPT1A* (Supp. Fig. 7a), cell viability was high (>90%), and there was no difference in diameter between cells exposed to oleic acid and control (Supp. Fig. 7b-e).

**FIGURE 3.**
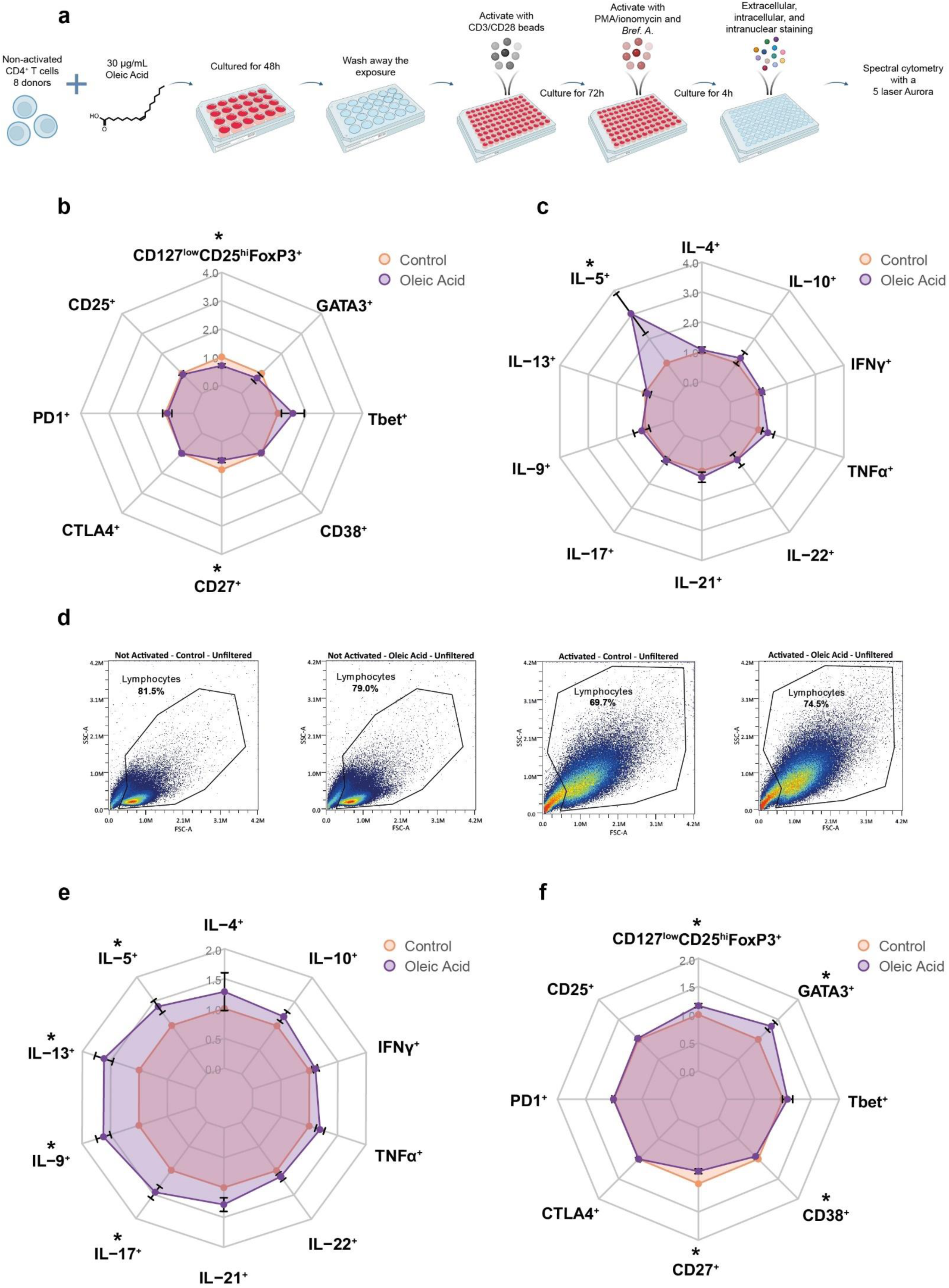
Oleic acid pre-exposure leads to changes in expression of extracellular markers, transcription factors, and intracellular cytokines. (*) PFDR < 0.05, n = 8. **(a)** Experimental set up for spectral cytometry measurements of oleic acid exposed non-activated CD4^+^ T cells for 48h with and without activation for 72h post-exposure. **(b)** Radar plot of various CD4^+^ T cell external markers and transcription factors expressed in CD4^+^ T cells after 48h of oleic acid exposure or control followed by 72h of rest and 4h stimulus with PMA/ionomycin. Values are expressed as fold change and standard error relative to control. **(c)** Radar plot of various CD4^+^ T cell internal cytokines expressed in CD4^+^ T cells after 48h of oleic acid exposure or control followed by 72h of rest and 4h stimulus with PMA/ionomycin. Values are expressed as fold change and standard error relative to control. **(d)** Forward and side scatter of activated vs. non-activated and control vs. oleic acid pre-exposed CD4^+^ T cells. Large differences in cell shape between the non-activated and activated state were observed but little difference in cell shape between pre-exposure to control or oleic acid were found. Non-activated control exposed cells are on the far left, non-activated oleic acid exposed cells are on the center left, activated control exposed cells are on the center right and activated oleic acid exposed cells are on the far right. **(e)** Radar plot of various CD4^+^ T cell internal cytokines expressed in CD4^+^ T cells after 48h of oleic acid exposure or control followed by 72h of activation with CD3/CD28 activation beads and 4h additional stimulus with PMA/ionomycin. Values are expressed as fold change and standard error relative to control. **(f)** Radar plot of various CD4^+^ T cell external markers and transcription factors expressed in CD4^+^ T cells after 48h of oleic acid exposure or control followed by 72h of activation with CD3/CD28 activation beads and 4h additional stimulus with PMA/ionomycin. Values are expressed as fold change and standard error relative to control.

First, we examined phenotypes after oleic acid exposure without activation (Supp. Table. 1g). We observed decreased frequencies of CD127^low^CD25^hi^FoxP3^+^ and CD27^+^ CD4^+^ T cells in response to oleic acid pre-exposure (P_FDR_ < 0.05; Fig. 3b). The CD127^low^CD25^hi^FoxP3^+^ population is representative of T_reg_ cells and thus, the decreased frequencies in the non-activated cells is in line with the lower *FOXP3* expression observed in the RNA sequencing analysis. Increased frequencies of IL-5^+^ cells were also observed (P_FDR_ < 0.05; Fig. 3c) These data suggest that the oleic acid-induced changes in gene expression are reflected in consistent functional characteristics of the CD4^+^ T cells without activation.

Activation of the CD4^+^ T cells led to an increased cell size irrespective of pre-exposure to oleic acid (Fig. 3d). In contrast, the expression of surface and intracellular markers was influenced by exposure to oleic acid prior to activation (Supp. Table. 1h). Pre-exposure to oleic acid resulted in a higher proportion of IL-9^+^ cells (P_FDR_ < 0.01) as compared to the control (Fig. 3e). Additional analysis showed that IL-9 was not co-expressed with other T_H_2 associated cytokines (Supp. Table. 1i). This aligns with our finding that a large percentage of upregulated genes mapped to a PU.1 motif (Fig. 2d), the key transcription factor controlling T_H_9 differentiation. Furthermore, increased frequencies of IL-17A^+^ cells were observed after pre-exposure to oleic acid as compared with control conditions (P_FDR_ < 0.05). As IL-17 is mainly produced by T_H_17 cells, it was hypothesized that other T_H_17 associated cytokines, such as IL-21, may also have been upregulated. Indeed, IL-21^+^ cells were increased in frequency (P < 0.05), but this effect was no longer significant after correction for multiple testing (P_FDR_ < 0.08). This aligns with our finding that a large percentage of upregulated genes mapped to the BHLHE40 motif (Fig. 2d), involved in T_H_17 differentiation^44, 45^. Activated CD4^+^ T cells showed increased frequencies of GATA3^+^ and CD127^low^CD25^hi^FoxP3^+^ and decreased frequencies of CD27^+^ and CD38^+^ cells in response to oleic acid pre-exposure (P_FDR_ < 0.05; Fig. 3f). GATA3 is the key transcription factor involved in T_H_2 differentiation and as such, frequencies of T_H_2-related cytokines IL-5^+^ and IL-13^+^ were increased (P_FDR_ < 0.05; Fig. 3e). Finally, we observed that the effect of oleic acid on differentiation is not secondary to a differential proliferative capacity (P > 0.92; Supp. Fig. 8). Together, these data indicate that the metabolic changes in non-activated CD4^+^ cells upon oleic acid exposure skew the cells towards producing more cytokines characteristic of T_H_9, T_H_17 and T_H_2 subsets upon activation.

### Oleic acid induced CD4^+^ T cell phenotypes blocked by metabolic inhibitors

We next determined whether induction of this profile, reminiscent of an increase differentiation towards T_H_9, T_H_17, and T_H_2 subsets was dependent on an upregulation of cholesterol and fatty acid biosynthesis in line with our RNA-seq data. We inhibited cholesterol synthesis with atorvastatin, targeting HMG-CoA reductase (*HMGCR*), and fatty acid synthesis with CP-640186, targeting both ACC1 and ACC2 (*ACACA* and *ACACB*). To this end, non-activated CD4^+^ T cells of 3 out of 8 donors for whom sufficient cells were available, were again exposed to control conditions, oleic acid only, oleic acid + 10μM atorvastatin, oleic acid + 20μM CP-640186, or oleic acid and both atorvastatin and CP-640186 for 48h. The effect of oleic acid exposure was confirmed by an upregulation of *CPT1A* (Supp. Fig. 9a). Cell viability was high (>88%) and there was no difference in diameter between cells exposed to control, oleic acid, or oleic acid + inhibitors (Supp. Fig. 9b-e).

Subsequently, both oleic acid and the inhibitors were washed away and the pre-exposed CD4^+^ T cells were activated. We evaluated the expression of one key marker for each subset: IL-9 for T_H_9, IL-17A for T_H_17, and IL-13 for T_H_2 cells (Fig. 4a). Remarkably, the ability of oleic acid to increase frequencies of IL-9^+^ cells was inhibited by both atorvastatin and CP-640186 (Fig. 4b: Supp. Table 1j). Although similar trends were observed for frequencies of IL-17A^+^ and IL-13^+^ cells, these effects were not statistically significant (Fig. 4b). These data indicate that oleic acid promotes the differentiation to in particular IL-9^+^ T cells via upregulation of cholesterol and fatty acid biosynthesis.

**FIGURE 4.**
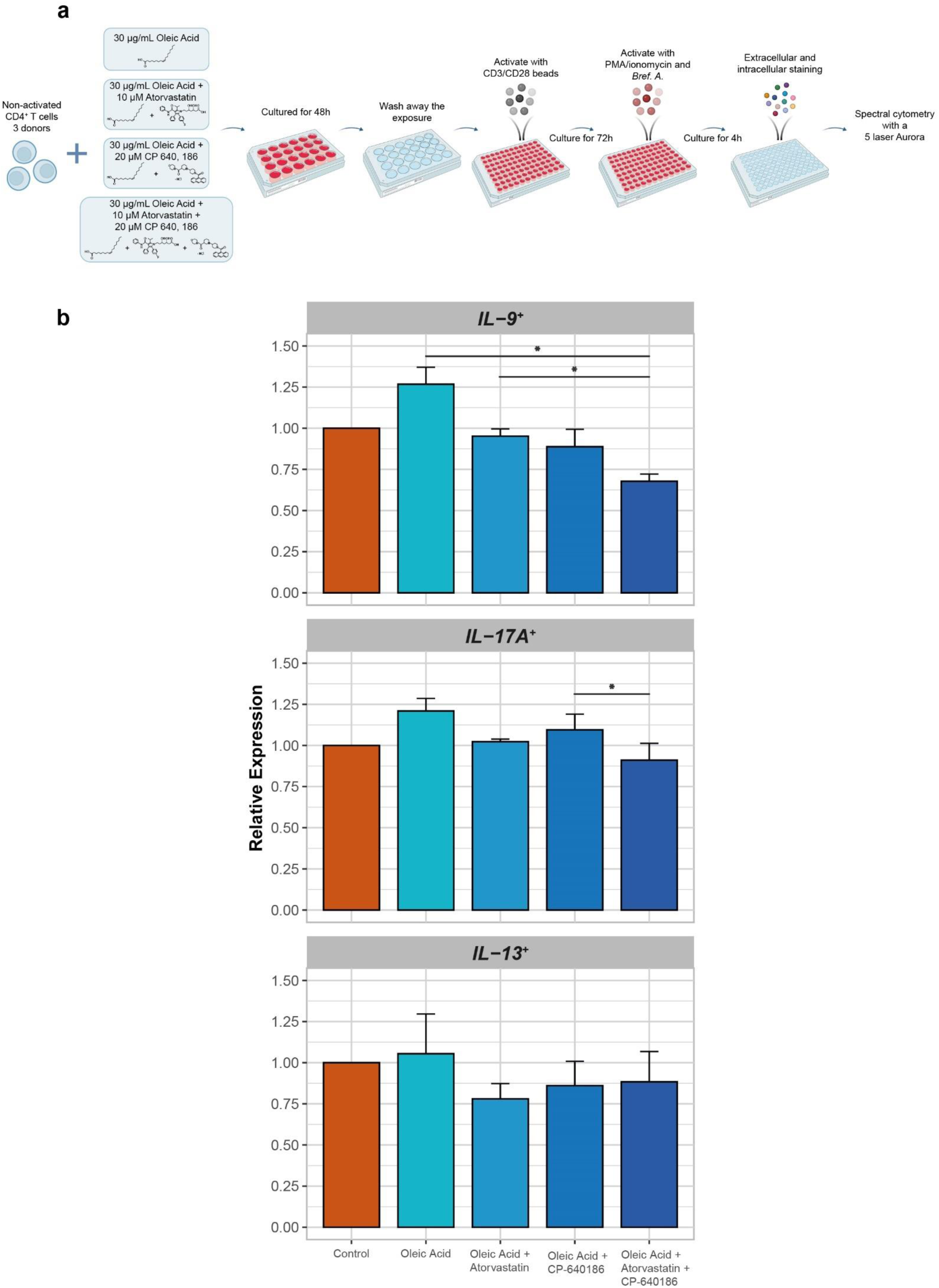
Metabolic inhibitors prevent oleic acid pre-exposure induced changes in expression of IL-9, IL-17A, and IL-13. (*) P < 0.05, n = 3. **(a)** Experimental set up for spectral cytometry measurements of oleic acid + inhibitor exposed non-activated CD4^+^ T cells for 48h with activation for 72h post-exposure. **(b)** Bar plot of IL-9, IL-17A, and IL-13 expression in CD4^+^ T cell after 48h of control, oleic acid, oleic acid + atorvastatin, oleic acid + CP-640186, or oleic acid + atorvastatin + CP-640186 exposure followed by 72h of activation with CD3/CD28 activation beads and 4h additional stimulus with PMA/ionomycin. Values are expressed as fold change and standard error relative to control.

## Discussion

T cells are known to respond to fatty acids^2^. Using an *in vitro* model of oleic acid exposure, we show that CD4^+^ T cells do so already when in a non-activated state by upregulating the expression of genes that encode enzymes involved in core metabolic pathways responsible for cholesterol biosynthesis, fatty acid biosynthesis, and aerobic glycolysis. These metabolic processes are hallmarks of activated and pro-inflammatory T cells^48^. Indeed, upon activation, CD4^+^ T cells pre-exposed to oleic acid are characterized by increased production of pro-inflammatory cytokines, including IL-9, IL-17A, IL-5, and IL-13, indicative of a preferential differentiation towards the pro-inflammatory T helper subsets T_H_9 and T_H_17 as well as T_H_2, which can have both pro- and anti-inflammatory effects. Interestingly, this effect is abolished in particular for T_H_9 cells by blocking the cholesterol or the fatty acid biosynthesis pathways during the initial exposure to oleic acid. Our findings imply that increased fatty acid levels in the circulation can rewire the metabolism of non-activated T cells and poise them to differentiate into more pro-inflammatory subsets for example when the cells infiltrate diseased tissues, including atherosclerotic plaques, and become activated.

Our results showed that cholesterol biosynthesis was the primary transcriptionally upregulated pathway in oleic acid exposed non-activated CD4^+^ T cells (15 out of 20 genes). This upregulation is of particular interest because of this pathway’s role in producing the necessary metabolites required for T cell activation^49^. Cholesterol biosynthesis is upregulated in activated T cells to support membrane production, cell signaling through the formation of lipid rafts, and prenylation of signaling proteins^50^. Additionally, intracellular cholesterol sensing has also been found to play a role in T cell differentiation, particularly towards pro-inflammatory subsets. For example, sterols were found to bind the T_H_17 transcription factor RORγt and could promote its activity^51^. Thus, the upregulation of gene expression in the cholesterol biosynthesis pathway due to oleic acid exposure may be indicative of a metabolic reprogramming of the non-activated CD4^+^ T cells towards an activated state and may lead to the differentiation towards pro-inflammatory subsets post-activation.

Additionally, expression of the two genes comprising the *de novo* fatty acid biosynthesis pathway were upregulated (*ACACA* and *FASN*). Together, cholesterol and fatty acid biosynthesis comprise part of the process known as lipogenesis, the synthesis of novel lipids in a cell. Lipogenesis is induced by the activation of the transcription factor SREBP1, which was associated to the upregulated transcripts in our RNA sequencing data. Enrichment analysis of our transcripts also revealed upregulated genes in mTORC1 signaling, which is known to induce the activation of SREBP1^52^. Although this effect is usually insulin dependent, obesity and overfeeding have been shown to hyperactivate mTORC1^53^. Thus, it is possible that oleic acid alone could induce the activation of mTORC1, which in turn activates SREBP1, leading to lipogenesis and expression of cholesterol and fatty acid biosynthesis related genes.

Fatty acid biosynthesis has also been related to the development of T_H_17 cells^17, 32^. Specifically, the mRNA expression of genes *ACACA*, encoding for acetyl-CoA carboxylase 1 (ACC1) and *FASN*, encoding fatty acid synthase, were increased in our dataset. These genes are key determinants in the development of the pro-inflammatory subset T_H_17 cells over the anti-inflammatory subset T_reg_ cells^22, 31, 42, 54, 55^. Correspondingly, *FOXP3*, the key transcription factor of T_reg_ cells, was downregulated in oleic acid exposed non-activated CD4^+^ T cells. Upregulated transcripts were found to be associated with the transcription factor PU.1. PU.1 is the key transcription factor in the development of T_H_9 cells. This subset is a highly pro-inflammatory subset related to T_H_2 cells^56^. This further supports the idea that oleic acid exposure leads to a cellular metabolic reprogramming that could promote the development of pro-inflammatory T cell subsets, specifically T_H_9, T_H_17, and possibly also T_H_2 cells. However, this effect was not dependent on epigenetic reprogramming as we measured no changes in DNA methylation. These results indicate that oleic acid-exposed non-activated CD4^+^ T cells were upregulating genes involved in metabolism to initiate/prepare for the selective differentiation into T_H_2/T_H_9/T_H_17 cells post-activation. Moreover, the metabolic processes being enhanced due to oleic acid exposure hint that the cells may preferentially differentiate towards T_H_2, T_H_9, and T_H_17 cells upon activation.

Importantly, we provide evidence that the oleic acid-induced metabolic rewiring underpins the observed enhanced T_H_9, T_H_17, and T_H_2 differentiation as exposing non-activated CD4^+^ T cells to oleic acid in combination with cholesterol or fatty acid synthesis inhibitors decreased the frequencies of IL-9^+^, IL-17A^+^, and IL-13^+^ cells. While the role of T_H_17 and T_H_2 cells in atherosclerosis has not been resolved, these cell types have been identified as pro-inflammatory in other diseases such as autoimmune encephalomyelitis and allergy, respectively^35, 57^. In contrast, T_H_9 cells have been implicated in atherosclerosis pathogenesis^58–60^.

Immune-lipid interactions occur in the circulation, which is a complex environment comprised of many factors that can affect T cell function prior to their recruitment to disease sites like the atherosclerotic plaque^61^. Fatty acids are a significant component of this environment and have been found to exert their effect not only on atherosclerosis but also on T cell function^2^. Oleic acid is one of the most abundant fatty acids in the human circulation^36^ and has been found to have pro-atherogenic effects^37^. However, our study does not preclude that other fatty acids, under polarizing conditions or *in vivo*, also affect non-activated CD4^+^ T cells and promote pro- or anti-inflammatory effects relevant for disease. Additionally, statins have been doted as having protective effects independent of cholesterol reduction^62^, our study hints that statins effect on T cell responses could contribute to this protective role.

Taken together, our results suggest that oleic acid can rewire the metabolism of non-activated CD4^+^ T cells, as they exist in circulation. Furthermore, this metabolic rewiring induces a preferential differentiation toward pro-inflammatory T cell types following activation, which is blocked by inhibiting cholesterol biosynthesis and fatty acid biosynthesis. Diseased sites, such as the atherosclerotic plaque, may induce differentiation into such proinflammatory subsets and this process may contribute to the atherogenic effect observed of CD4^+^ T cells^7–9^.

## Acknowledgements

The authors’ work is supported by the Dutch CardioVascular Alliance (The Dutch Heart Foundation, Dutch Federation of University Medical Centers, the Netherlands Organization for Health Research and Development, and the Royal Netherlands Academy of Sciences) for the GENIUSI and GENIUSII projects Generating the Best Evidence-Based Pharmaceutical Targets for Atherosclerosis (CVON2011-19, CVON2017-20, respectively) and the Joint Programming Initiative a Healthy Diet for a Healthy Life (JPI HDHL) administered by ZonMW, the Netherlands (grant 529051021).

## Author Contributions

B.T.H and J.W.J conceived the project. N.A.R. designed and conducted the experiments, analyzed the results, and drafted the manuscript. F.S. designed, and analyzed the spectral flow cytometry experiments. K.F.D. designed the *in vitro* model and analyzed the RNA sequencing data. J.C.K. and H.M.S. designed of the *in vitro* model. L.C. analyzed the DNA methylation data. S.H. designed the functional assays. M.A.H. performed and analyzed the transcription factor footprint analysis. H.M. aligned the RNA sequencing data. E.W.Z. conceived the statistical model used in the analysis of the RNA sequencing and DNA methylation data. B.E. conceived and interpreted the functional assays and spectral flow cytometry data. A.I.F. conceived and designed the *in vitro* model. All authors contributed to the writing of the manuscript.

## Methods

### Peripheral blood CD4^+^ T cell isolation and culture conditions

To obtain non-activated CD4^+^ T cells, peripheral blood mononuclear cells (PBMCs) were isolated from buffy coats of anonymous blood bank donors (Sanquin, Amsterdam, The Netherlands) by Ficoll paque (Apotheek LUMC, 97902861) gradient centrifugation. Next, CD4^+^ T cells were purified from the PBMCs using lyophilized human anti-CD4^+^ magnetically labeled microbeads (Miltenyi, 130-097-048) scaling the manufacturer’s instructions to ⅕ of the recommended volumes. CD4^+^ T cell purity was assessed on an LSR-II instrument at the Leiden University Medical Center Flow Cytometry Core Facility (https://www.lumc.nl/research/facilities/fcf/) with the BD FACSDiva™ v8.0.2 software (BD Biosciences). Cells were stained with anti-CD3-PE (BD Biosciences, 345765), anti-CD4-APC (BD Biosciences, 345771), anti-CD8-FITC (BD Biosciences, 555634), and anti-CD14-PEcy7 (BD Biosciences, 557742) and resuspended in 1% paraformaldehyde (Apotheek LUMC, 120810-001) to fix the cells prior to acquisition. Purity was >98% for all donors.

Prior to oleic acid exposure, ∼8*10^7^ isolated cells were cultured overnight to allow the cells to return to a resting state after the stress of the isolation procedure. This was done in T75 flasks (Greiner Bio-One, 658-175) at a density of ∼2.5*10^6^ cells/mL in 5% fetal calf serum (FCS) (Bodinco BDC, 16941) DMEM (Dulbecco’s Modified Eagle’s Serum (Sigma, 05796), 1% Pen-Strep (Lonza, DE17-602E), 1% GlutaMAX-1 (100x) (Gibco, 35050-038)) medium supplemented with 50 IU/mL IL-2 (Peprotech, 200-02) and incubated at 37°C under 5% CO_2_. To keep the cells in a non-activated state, no additional stimulus was added. Any CD4^+^ T cells not used directly after the isolation were kept in DMEM supplemented with 30% FCS, 1% Pen-Strep, 1% GlutaMAX-1, and 20% Dimethyl Sulfoxide (DMSO) (WAK-Chemie Medical GmbH, WAK-DMSO-10) medium, and stored in liquid nitrogen.

Next, non-activated CD4^+^ T cells were cultured with or without oleic acid for 0.5, 3, 24, 48, or 72 hours at 37°C under 5% CO_2_. To this end, CD4^+^ T cells from each donor were plated in a 24 wells plate (density of ∼4*10^6^ cells/well) in 2mL 5% FCS DMEM for each time point, one exposed to oleic acid, one to the solvent control, and one to the negative control (Fig. 1A). Cells were cultured in medium containing FCS to ensure cell viability during culture and to be more comparable to physiological conditions of the circulation where other lipids are also present. Oleic acid (Sigma, O1383) was dissolved in HPLC grade ethanol (Fisher Scientific, 64-17-5) to a final concentration of 30,000μg/mL and complexed to fatty acid-free (FAF) bovine serum albumin (BSA) (Sigma, A7030) in a 2% FAF BSA DMEM mixture (Dulbecco’s Modified Eagle’s Serum, 2% FAF BSA, 1% Pen-Strep, 1% GlutaMAX-1 (100x)) to a final concentration of 150μg/mL. Complexing oleic acid mimics physiological conditions as fatty acids are also bound to albumin in the human circulation^63^. Oleic acid was further diluted to the final concentrations of 10, 20, 30, and 50μg/mL. The concentrations tested were determined based on a literature search^10–12, 38–42^. For the solvent control samples, HPLC grade ethanol was diluted in 2% FAF BSA DMEM in the same volume as to dilute oleic acid to 150μg/mL and added to the wells. For the negative control samples, 2% FAF BSA DMEM was added directly to the wells with no additional solvent. The amount of 2% FAF BSA DMEM added to the wells was equal for each condition to keep the volumes equivalent. To assess the additional oleic acid stimulus to the non-activated CD4^+^ T cells due to FCS in the culture medium, an FCS sample was measured via the Shotgun Lipidomics Assistant (SLA) method^64^ to estimate the fraction of oleic acid in the sample. The sample was prepped as previously described^65^ but with two modifications, a starting volume of 25μL FCS and 600μL MTBE was added instead of 575μL during the first extraction. After exposure, the cells were flash frozen in liquid nitrogen and stored at -80°C until further use. Cell viability was measured via trypan blue staining (Sigma Aldrich, T8154).

### RNA and DNA Isolation

To isolate total RNA for RNA sequencing and RT-qPCR and genomic DNA for 850k EPIC DNA methylation array, RNA and DNA were simultaneously extracted from the cell samples using the Zymo Quick-DNA/RNA Microprep Plus Kit (Zymo Research, D7005) according to manufacturer’s instructions. The RNA was quantified using a Qubit® 2.0 Fluorometer (Q32866) with the Qubit® RNA BR Assay Kit (Thermofisher, Q10211) according to manufacturer’s instructions. RNA integrity (RIN) values of the samples were on average 8.40 SE 0.14 as determined using an Agilent 2100 Bioanalyzer Instrument (G2939BA) with the Agilent RNA 6000 Nano Reagents (Agilent, 5067-1511). The DNA was quantified using a Qubit® 2.0 Fluorometer (Q32866) with the Qubit® DNA HS Assay Kit (Thermofisher, Q10211) according to manufacturer’s instructions. RNA was divided into two samples and stored at -80°C, 1μg for RNA sequencing and the rest for cDNA synthesis and RT-qPCR measurements. DNA was stored at -80°C, for 850k EPIC DNA methylation measurements.

### Real time-quantitative PCR

To measure the expression of *CPT1A* in all the cell samples, cDNA was synthesized with 200ng of the stored RNA using the Transcriptor First Strand cDNA Synthesis Kit (Roche, 04897030001) according to the manufacturer’s instructions. Quantitative real time PCR’s for *CPT1A* (Thermofisher, Hs00912671_m1, 4331182) were performed using the TaqMan™ Fast Advanced Master Mix (Thermofisher, 4444557) with 10ng cDNA per reaction on a QuantStudio 6 Real-Time PCR system (Applied Biosystems). All RT-qPCR reactions were performed in triplicate and outliers were removed if the Ct value measured differed more than 0.5% from the mean. Relative gene expression levels (-ΔCt) were calculated using the average of Ct values of *RPL13A* (Thermofisher, Hs03043887_gH, 4448892) and *SDHA* (Thermofisher, Hs00188166_m1, 4453320) as internal controls^66^. The fold change was determined using the 2^-ΔΔCt^ method, using the negative control as the reference. All statistical analyses were performed in R. Data are expressed as mean of the relative fold change and standard error. The reported P values were determined by applying a paired two-tailed student’s T test. P values < 0.05 were considered to be statistically significant.

### RNA Sequencing Analysis

RNA sequencing (RNA-seq) was performed to determine the differences in the transcriptome of oleic acid versus solvent exposed non-activated CD4^+^ T cells across time. 1μg of total RNA from each of the samples was sent for sequencing (Macrogen, Amsterdam, NL), each with a concentration above 20ng/μL in 50μL solution. RNA-seq libraries were prepared from 200ng RNA using the Illumina Truseq stranded mRNA library prep (Illumina, 20020595) after depletion of ribosomal RNA with Ribo Zero Gold (Illumina, 20037135). Both whole-transcriptome amplification and sequencing library preparations were performed in two 96-well plates with half the samples each, to reduce assay-to-assay variability. Quality control steps were included to determine total RNA quality and quantity, the optimal number of PCR preamplification cycles, and fragment size selection. No samples were eliminated from further downstream steps. Barcoded libraries were pooled and equally divided across two lanes to ensure an equal distribution of all the samples across the two lanes. Barcoded libraries were sequenced to a read depth of 30 million reads using the Novaseq 6000 (Illumina) to generate 100 base pair paired-end reads.

RNA-seq reads were processed using the BioWDL RNAseq pipeline (v3.0.0) developed at LUMC (http://zenodo.org/record/3713261#.ZF98HdJBw5k). Quality controls were performed using FastQC (v0.11.7) and MultiQC (v1.7). Cleaned reads were aligned to the human reference genome GRCh38 using STAR aligner (v2.7.3a). Gene count table was generated using Htseq-count (v0.11.2) with Ensembl gene annotation version 99. Based on Ensembl gene biotype annotation, we included only protein coding genes for further downstream analysis (19,916 genes in total). We used the Bioconductor package *DESeq2*^67^ (v1.40.1) to test whether oleic acid had an effect on gene expression at any time point. DESeq fits a generalized linear model (GLM) assuming the negative binomial distribution for the counts. The model expresses the logarithm of the average of the counts in terms of one of more predictors. In this case, we compared two models: The first “null” model has only timepoint (as a categorical variable with 5 levels) and subject identifier as predictors. By including the subject identifier in the model, we account for the dependence between measurements within the same subject. The second “alternative” model also includes the interaction between phenotype (oleic acid as a numerical measurement) and timepoint. We compare the fit of the two models with a likelihood ratio test. As part of the DESeq2 process lowly expressed genes were automatically removed, resulting in 12,932 analyzed genes^67^. The Benjamini-Hochberg procedure was used to correct for multiple testing and a false discovery rate (FDR) < 0.05 was considered statistically significant.

Next, to identify distinct gene expression patterns in the data, unsupervised K means clustering was performed on the differentially expressed genes using the *factoextra*^68^ package (v1.0.7). The number of clusters, *k*, was chosen using the elbow, silhouette, and gap-statistic method. Heatmaps were constructed using *ComplexHeatmap*^69^ (v2.14.0) by plotting the log2FoldChange of the DEGs at each time point (Supp. Table 1a and b).

The identified clusters were then mapped for pathway enrichment. 10 human pathway databases (BioPlanet 2019, WikiPathways 2019 Human, KEGG 2019 Human, Elsevier Pathway Collection, BioCarta 2015, Reactome 2016, HumanCyc 2016, NCI-Nature 2016, Panther 2016 and MSigDB Hallmark 2020) were queried using gene symbols, with 430 of 544 queried genes present in at least 1 database. The identified clusters were then mapped for pathway enrichment using *clusterProfiler*^70^ (v4.6.2) with the background set to 12,932 expressed genes in the CD4^+^ T cells based on *DESeq2* filtering. Multiple testing using the Benjamini-Hochberg method at 5% FDR was performed over the combined results from the 10 databases. Pathways that included highly similar gene sets were grouped (Jaccard index > 0.7) and only the most significantly enriched pathway per group was retained. Furthermore, using the UniProt IDs of the enriched genes (Supp. Table 1a and b), the Path-MAP function of the PathBank database^71^ was used to visualize the list of matching components within specific canonical pathways. *De novo* motif analysis on promotors of differentially regulated genes was performed using HOMER^72^.

### DNA methylation analysis

To investigate if DNA methylation of non-activated CD4^+^ T cells was affected by oleic acid exposure, genome-wide DNA methylation was assessed using Illumina EPIC DNA methylation array. DNA was diluted to a final concentration of >500ng in 45μL buffer. Samples were randomized for exposure, timepoint, and donor ID, using the *Omixer*^73^ (v4.3) package in R, across two 96-well plates with 56 samples on one plate and 34 on the other, to minimize batch effects. Samples were then sent for Illumina 850k EPIC DNA methylation array measurements (Research Unit Molecular Epidemiology, Helmholtz Institute, Munich, DE). Here, genomic DNA was bisulfite converted using the EZ-96 DNA Methylation Kit (Zymo Research, D5004) according to the manufacturer’s instructions. Subsequent methylation analysis was performed on an Illumina (San Diego, CA, USA) iScan platform using the Infinium MethylationEPIC BeadChip according to standard protocols provided by Illumina. Briefly, a whole genome amplification step was performed using 4µL of each bisulfite converted sample, followed by enzymatic fragmentation and application of the samples to BeadChips (Illumina). The arrays were fluorescently stained and scanned. GenomeStudio software (v2011.1) with Methylation Module (v1.9.0) was used for initial quality control of assay performance and for generation of methylation data export files.

Following receipt of the methylation data as IDAT files, preprocessing and quality control (QC) followed the DNAmArray pipeline^74^. Briefly, sample quality was assessed using visualizations, including *MethylAid*^75^ (v1.32.0) plots. The data underwent functional normalization using 6 principal components, followed by removal of outlying or unreliable values, including those based on low bead number (0.16%), intensity (0.03%), or that were not substantially distinguishable from background noise (0.42%). Additionally, cross-reactive and polymorphic^76^ probes were omitted, along with those on sex chromosomes. The resulting dataset contained DNA methylation data at 738,991 CpGs for 90 samples and was annotated using the IlluminaHumanMethylationEPIC manifest (v1.0b2). Linear models were fitted for each probe using the *limma*^77^ package (v3.54.2) in R, with DNA methylation as the dependent and stimulation status as the independent variable. Timepoint and donor ID were adjusted for, and the effect of stimulation status was allowed to vary by timepoint using an interaction term. P-values were adjusted for multiple testing using FDR, with a 5% significance threshold.

### Culture Conditions and T cell Activation

To study the effect of oleic acid pre-exposure on CD4^+^ T cell subset development, cells from 8 out of 9 donors that were previously analyzed using RNA-seq were thawed from liquid nitrogen; 1 donor could not be studied further because too few cells were available. Cells were cultured overnight to allow the cells to return to a resting state after the stress of the thawing, in T75 flasks at a density of ∼2.5*10^6^ cells/mL in 5% FCS DMEM medium supplemented with 50 IU/mL IL-2 at 37°C under 5% CO_2_. To keep the cells in a non-activated state, no additional stimulus was added. Following overnight incubation, the cells were divided into 2 conditions, oleic acid and solvent exposed, and plated in a 24 wells plate (density of ∼4*10^6^ cells/well) in 2mL 5% FCS DMEM. The oleic acid and solvent solution were prepared as stated previously, with one modification. To ensure that there was no effect of the solvent on T cell differentiation, the HPLC grade EtOH was evaporated before dissolving the oleic acid in 2% FAF BSA DMEM medium. The HPLC grade EtOH was also evaporated before adding the 2% FAF BSA DMEM medium in the solvent exposed condition, rendering it essentially the same as the negative control. These solutions were each added to the respective wells, where the final concentration of the oleic acid exposed conditions equaled 30μg/mL. The CD4^+^ T cells were cultured for 48h at 37°C under 5% CO_2_.

To ensure that the effect on CD4^+^ T cell differentiation was due to oleic acid pre-exposure, all medium of each condition was replaced by 5% FCS medium after 48h of exposure, before initiating the activation. Cell viability and diameter were first measured by Via1-Cassette™ (Chemometec, 941-0012) on a NucleoCounter® NC-200™ (Chemometec, 900-0200) and found to be > 90% for each condition. Then, 2 million cells were harvested by flash freezing in liquid nitrogen for *in vitro* model confirmation by RT-qPCR. The remaining cells were plated in a round bottom 96 wells plate (Corning Incorporated, 3799), at a density of 100,000 cells/well, and were activated for 72h using Dynabeads™ Human T-Activator CD3/CD28 for T Cell Expansion and Activation (Thermofisher, 11161D) according to the manufacturer’s instructions at 37°C under 5% CO_2_. Half the cells from each exposure were activated and the other half was left in the non-activated state. Subsequently, the cells from each pre-exposure and activation state were pooled in Eppendorf tubes and the beads were magnetically removed from the activated cells. Cell viability and diameter were measured by Via1-Cassette™ after 72h. Cells were then used for T cell subset identification described in more detail below. All centrifugation steps were performed at 1500 rpm at room temperature.

### Inhibitor Culture Conditions and T cell Activation

To study whether the effect of oleic acid pre-exposure on CD4^+^ T cell subset development could be prevented by metabolic inhibitors, cells from 3 out of 8 donors that were previously analyzed for subset development were thawed from liquid nitrogen; 5 donors could not be studied further because too few cells were available. Cells were cultured overnight to allow the cells to return to a resting state after the stress of the thawing, in T75 flasks at a density of ∼2.5*10^6^ cells/mL in 5% FCS DMEM medium supplemented with 50 IU/mL IL-2 at 37°C under 5% CO_2_. To keep the cells in a non-activated state, no additional stimulus was added. Following overnight incubation, the cells were divided into 5 conditions, solvent, oleic acid, oleic acid + atorvastatin (Sigma Aldrich, PHR1422), oleic acid + CP- 640186 (Sanbio, 17691-5), and oleic acid + atorvastatin + CP-640186 exposed, and plated in a 24 wells plate (density of ∼4*10^6^ cells/well) in 2mL 5% FCS DMEM. The oleic acid and solvent solution were prepared as stated previously, with HPLC grade EtOH evaporation. These solutions were each added to the respective wells, where the final concentration of the oleic acid exposed conditions equaled 30μg/mL. Atorvastatin and CP-640186 were added to the respective wells at a concentration of 10μM and 20μM, respectively. The CD4^+^ T cells were cultured for 48h at 37°C under 5% CO_2_.

To ensure that the effect on CD4^+^ T cell differentiation was due to oleic acid and inhibitor pre-exposure, all medium of each condition was replaced by 5% FCS medium after 48h of exposure, before initiating the activation. Cell viability and diameter were first measured by Via1-Cassette™ (Chemometec, 941-0012) on a NucleoCounter® NC-200™ (Chemometec, 900-0200) and found to be > 90% for each condition. Then, ∼0.5-1.5 million cells were harvested by flash freezing in liquid nitrogen for *in vitro* model confirmation by RT-qPCR. The remaining cells were plated in a round bottom 96 wells plate (Corning Incorporated, 3799), at a density of 100,000 cells/well, and were activated for 72h using Dynabeads™ Human T-Activator CD3/CD28 for T Cell Expansion and Activation (Thermofisher, 11161D) according to the manufacturer’s instructions at 37°C under 5% CO_2_. Subsequently, the cells from each pre-exposure were pooled in Eppendorf tubes and the beads were magnetically removed. Cell viability and diameter were measured by Via1-Cassette™ after 72h. Cells were then used for T cell subset identification described in more detail below. All centrifugation steps were performed at 1500 rpm at room temperature.

### Spectral Cytometry

Prior to FACS analysis, cells were washed in RPMI 1640 medium (Thermofisher, 42401), supplemented with 100U/mL penicillin, 100µg/mL streptomycin, 1mM pyruvate, 2mM glutamate, and 10% FCS (Serana, S-FBS-SA-015), and adjusted to a concentration of 1x10^6^ cells/mL. Cells were then resuspended in 100µL RPMI + 10% FCS and stimulated for 4h with PMA (100ng/mL, Merck, P8139) and ionomycin (1µg/mL, Merck, I0634) at 37°C under 5% CO_2_ to promote cytokine production^78^. After 2h of stimulation, 10µg/mL of the protein transport inhibitor Brefeldin A (Merck, B7651) was added.

After stimulation, the cells were washed twice in phosphate-buffered saline (PBS), stained for viable cells with LIVE/DEAD™ Fixable Blue (Thermofisher, L34962) for 30min at room temperature, then washed twice in fluorescence-activated cell sorting (FACS) buffer (PBS supplemented with 0.5% BSA (Merck, 10735086001) and 2mM EDTA (Invitrogen, 15575020)). The antibody surface cocktail (11 markers, see Supp. Table 1k), prepared in FACS buffer containing 20% Brilliant Stain Buffer Plus (BD Biosciences, 563794) was added to the cells and incubated for 30min at room temperature. Cells were then washed twice in FACS buffer and afterwards fixed and permeabilized with the Fixation/Permeabilization solution from the eBioscience™ FoxP3 / Transcription Factor Staining Buffer Set (Thermofisher, 00-5523-00) according to the manufacturer’s instructions for 30min at 4°C. Subsequently, cells were washed twice with the Permeabilization buffer from the eBioscience™ FoxP3 / Transcription Factor Staining Buffer Set before being stained with the intracellular/intranuclear antibody cocktail for 30min at 4°C (14 markers, see Supp. Table 1k). Lastly, cells were washed with eBioscience™ Permeabilization buffer followed by another wash in FACS buffer. All centrifugation steps before fixation were performed at 300x g at room temperature and after fixation at 800x g at 4°C. Single-stain reference controls were either cells or UltraComp eBeads™ (Supp. Table 1k), while cells were used as unstained reference control (Supp. Table 1k). All reference controls underwent the same protocol as the fully stained samples, including washes, buffers used, and fixation and permeabilization steps.

For acquisition, cells were resuspended in FACS buffer and acquired on a 5L-Cytek Aurora instrument at the Leiden University Medical Center Flow Cytometry Core Facility with the SpectroFlo® v2.2.0.3 software (Cytek Biosciences). Data was manually gated in OMIQ (Dotmatics, 2023). All statistical analyses were performed in R. Data are expressed as mean of the relative fold change and standard error. The reported P values were determined by applying a paired two-tailed student’s T test. Differences with P_FDR_ < 0.05 (Benjamini-Hochberg) were considered to be significant.

## Data Availability

The data supporting the findings of this study are available within the article and its Supplementary information files. All other data including the raw files are available at the Gene Expression Omnibus repository, accession GEO (main combined submission: GSE231459, RNA sequencing submission: GSE231458, and DNA methylation submission: GSE231457).

## References

1. Schaftenaar, F., Frodermann, V., Kuiper, J. & Lutgens, E. Atherosclerosis: the interplay between lipids and immune cells. Curr Opin Lipidol 27, 209–215, (2016).

2. Reilly, N. A., Lutgens, E., Kuiper, J., Heijmans, B. T. & Wouter Jukema, J. Effects of fatty acids on T cell function: role in atherosclerosis. Nat Rev Cardiol 18, 824–837, (2021).

3. Winkels, H. et al. Atlas of the Immune Cell Repertoire in Mouse Atherosclerosis Defined by Single-Cell RNA-Sequencing and Mass Cytometry. Circ Res 122, 1675–1688, (2018).

4. Fernandez, D. M. et al. Single-cell immune landscape of human atherosclerotic plaques. Nat Med 25, 1576–1588, (2019).

5. Depuydt, M. A. C. et al. Microanatomy of the Human Atherosclerotic Plaque by Single-Cell Transcriptomics. Circ Res 127, 1437–1455, (2020).

6. Zernecke, A. et al. Meta-Analysis of Leukocyte Diversity in Atherosclerotic Mouse Aortas. Circ Res 127, 402–426, (2020).

7. Ketelhuth, D. F. & Hansson, G. K. Adaptive Response of T and B Cells in Atherosclerosis. Circ Res 118, 668–678, (2016).

8. Saigusa, R., Winkels, H. & Ley, K. T cell subsets and functions in atherosclerosis. Nat Rev Cardiol 17, 387–401, (2020).

9. Zhou, X., Robertson, A. K., Hjerpe, C. & Hansson, G. K. Adoptive transfer of CD4+ T cells reactive to modified low-density lipoprotein aggravates atherosclerosis. Arterioscler Thromb Vasc Biol 26, 864–870, (2006).

10. Angela, M. et al. Fatty acid metabolic reprogramming via mTOR-mediated inductions of PPARgamma directs early activation of T cells. Nat Commun 7, 13683, (2016).

11. Ioan-Facsinay, A. et al. Adipocyte-derived lipids modulate CD4+ T-cell function. Eur J Immunol 43, 1578–1587, (2013).

12. Hossein zade, A. Fatty Acids Effect on T Helper Differentiation in Vitro. International Journal of Nutrition and Food Sciences 5, (2016).

13. Bi, X. et al. omega-3 polyunsaturated fatty acids ameliorate type 1 diabetes and autoimmunity. J Clin Invest 127, 1757–1771, (2017).

14. Raphael, I., Nalawade, S., Eagar, T. N. & Forsthuber, T. G. T cell subsets and their signature cytokines in autoimmune and inflammatory diseases. Cytokine 74, 5–17, (2015).

15. Grivel, J. C. et al. Activation of T lymphocytes in atherosclerotic plaques. Arterioscler Thromb Vasc Biol 31, 2929–2937, (2011).

16. Geltink, R. I. K., Kyle, R. L. & Pearce, E. L. Unraveling the Complex Interplay Between T Cell Metabolism and Function. Annu Rev Immunol 36, 461–488, (2018).

17. Chapman, N. M., Boothby, M. R. & Chi, H. Metabolic coordination of T cell quiescence and activation. Nat Rev Immunol 20, 55–70, (2020).

18. MacIver, N. J., Michalek, R. D. & Rathmell, J. C. Metabolic regulation of T lymphocytes. Annu Rev Immunol 31, 259–283, (2013).

19. Warburg, O., Gawehn, K. & Geissler, A. W. Stoffwechsel der weissen Blutzellen [Metabolism of leukocytes]. Z Naturforsch B. 13B, 515-516, (1958).

20. Vander Heiden, M. G., Cantley, L. C. & Thompson, C. B. Understanding the Warburg Effect: The Metabolic Requirements of Cell Proliferation. Science 324, 1029–1033, (2009).

21. Howie, D., Ten Bokum, A., Necula, A. S., Cobbold, S. P. & Waldmann, H. The Role of Lipid Metabolism in T Lymphocyte Differentiation and Survival. Frontiers in Immunology 8, (2018).

22. Cluxton, D., Petrasca, A., Moran, B. & Fletcher, J. M. Differential Regulation of Human Treg and Th17 Cells by Fatty Acid Synthesis and Glycolysis. Front Immunol 10, 115, (2019).

23. Michalek, R. D. et al. Cutting edge: distinct glycolytic and lipid oxidative metabolic programs are essential for effector and regulatory CD4+ T cell subsets. J Immunol 186, 3299–3303, (2011).

24. Fullerton, M. D., Steinberg, G. R. & Schertzer, J. D. Immunometabolism of AMPK in insulin resistance and atherosclerosis. Mol Cell Endocrinol 366, 224–234, (2013).

25. Maganto-Garcia, E., Tarrio, M. L., Grabie, N., Bu, D. X. & Lichtman, A. H. Dynamic changes in regulatory T cells are linked to levels of diet-induced hypercholesterolemia. Circulation 124, 185–195, (2011).

26. Delgoffe, G. M. et al. The mTOR kinase differentially regulates effector and regulatory T cell lineage commitment. Immunity 30, 832–844, (2009).

27. Korn, T., Bettelli, E., Oukka, M. & Kuchroo, V. K. IL-17 and Th17 Cells. Annu Rev Immunol 27, 485–517, (2009).

28. Zhu, J., Yamane, H. & Paul, W. E. Differentiation of effector CD4 T cell populations (*). Annu Rev Immunol 28, 445–489, (2010).

29. Delgoffe, G. M. et al. The kinase mTOR regulates the differentiation of helper T cells through the selective activation of signaling by mTORC1 and mTORC2. Nat Immunol 12, 295–303, (2011).

30. Shi, L. Z. et al. HIF1alpha-dependent glycolytic pathway orchestrates a metabolic checkpoint for the differentiation of TH17 and Treg cells. J Exp Med 208, 1367–1376, (2011).

31. Young, K. E., Flaherty, S., Woodman, K. M., Sharma-Walia, N. & Reynolds, J. M. Fatty acid synthase regulates the pathogenicity of Th17 cells. J Leukoc Biol 102, 1229–1235, (2017).

32. Berod, L. et al. De novo fatty acid synthesis controls the fate between regulatory T and T helper 17 cells. Nat Med 20, 1327–1333, (2014).

33. O’Sullivan, D. & Pearce, E. L. Fatty acid synthesis tips the TH17-Treg cell balance. Nat Med 20, 1235–1236, (2014).

34. Gerriets, V. A. & Rathmell, J. C. Metabolic pathways in T cell fate and function. Trends Immunol 33, 168–173, (2012).

35. Nakayama, T. et al. Th2 Cells in Health and Disease. Annu Rev Immunol 35, 53–84, (2017).

36. Bicalho, B., David, F., Rumplel, K., Kindt, E. & Sandra, P. Creating a fatty acid methyl ester database for lipid profiling in a single drop of human blood using high resolution capillary gas chromatography and mass spectrometry. J Chromatogr A 1211, 120–128, (2008).

37. Steffen, B. T., Duprez, D., Szklo, M., Guan, W. & Tsai, M. Y. Circulating oleic acid levels are related to greater risks of cardiovascular events and all-cause mortality: The Multi-Ethnic Study of Atherosclerosis. J Clin Lipidol 12, 1404–1412, (2018).

38. Passos, M. E. et al. Differential effects of palmitoleic acid on human lymphocyte proliferation and function. Lipids Health Dis 15, 217, (2016).

39. Verlengia, R. et al. Effect of arachidonic acid on proliferation, cytokines production and pleiotropic genes expression in Jurkat cells--a comparison with oleic acid. Life Sci 73, 2939–2951, (2003).

40. Stentz, F. B. & Kitabchi, A. E. Palmitic acid-induced activation of human T-lymphocytes and aortic endothelial cells with production of insulin receptors, reactive oxygen species, cytokines, and lipid peroxidation. Biochem Biophys Res Commun 346, 721–726, (2006).

41. Gorjao, R., Cury-Boaventura, M. F., de Lima, T. M. & Curi, R. Regulation of human lymphocyte proliferation by fatty acids. Cell Biochem Funct 25, 305–315, (2007).

42. Endo, Y. et al. ACC1 determines memory potential of individual CD4(+) T cells by regulating de novo fatty acid biosynthesis. Nat Metab 1, 261–275, (2019).

43. Abdelmagid, S. A. et al. Comprehensive profiling of plasma fatty acid concentrations in young healthy Canadian adults. PLoS One 10, e0116195, (2015).

44. Lin, C. C. et al. IL-1-induced Bhlhe40 identifies pathogenic T helper cells in a model of autoimmune neuroinflammation. J Exp Med 213, 251–271, (2016).

45. Nechanitzky, R. et al. Cholinergic control of Th17 cell pathogenicity in experimental autoimmune encephalomyelitis. Cell Death Differ 30, 407–416, (2023).

46. Kidani, Y. et al. Sterol regulatory element-binding proteins are essential for the metabolic programming of effector T cells and adaptive immunity. Nat Immunol 14, 489–499, (2013).

47. Shin, H. J., Lee, J. B., Park, S. H., Chang, J. & Lee, C. W. T-bet expression is regulated by EGR1-mediated signaling in activated T cells. Clin Immunol 131, 385–394, (2009).

48. Cai, F., Jin, S. & Chen, G. The Effect of Lipid Metabolism on CD4(+) T Cells. Mediators Inflamm 2021, 6634532, (2021).

49. Smith-Garvin, J. E., Koretzky, G. A. & Jordan, M. S. T cell activation. Annu Rev Immunol 27, 591–619, (2009).

50. Shyer, J. A., Flavell, R. A. & Bailis, W. Metabolic signaling in T cells. Cell Res 30, 649–659, (2020).

51. Santori, F. R. et al. Identification of natural RORgamma ligands that regulate the development of lymphoid cells. Cell Metab 21, 286–298, (2015).

52. Laplante, M. & Sabatini, D. M. An emerging role of mTOR in lipid biosynthesis. Curr Biol 19, R1046–1052, (2009).

53. Khamzina, L., Veilleux, A., Bergeron, S. & Marette, A. Increased activation of the mammalian target of rapamycin pathway in liver and skeletal muscle of obese rats: possible involvement in obesity-linked insulin resistance. Endocrinology 146, 1473–1481, (2005).

54. Endo, Y. et al. Obesity Drives Th17 Cell Differentiation by Inducing the Lipid Metabolic Kinase, ACC1. Cell Rep 12, 1042–1055, (2015).

55. Sun, L., Fu, J. & Zhou, Y. Metabolism Controls the Balance of Th17/T-Regulatory Cells. Front Immunol 8, 1632, (2017).

56. Angkasekwinai, P. & Dong, C. IL-9-producing T cells: potential players in allergy and cancer. Nat Rev Immunol 21, 37–48, (2021).

57. Schnell, A., Littman, D. R. & Kuchroo, V. K. T(H)17 cell heterogeneity and its role in tissue inflammation. Nat Immunol 24, 19–29, (2023).

58. Zhang, W. et al. IL-9 aggravates the development of atherosclerosis in ApoE-/- mice. Cardiovasc Res 106, 453–464, (2015).

59. Gregersen, I. et al. Increased systemic and local interleukin 9 levels in patients with carotid and coronary atherosclerosis. PLoS One 8, e72769, (2013).

60. Li, Q. et al. Increased Th9 cells and IL-9 levels accelerate disease progression in experimental atherosclerosis. Am J Transl Res 9, 1335–1343, (2017).

61. Visscher, M. et al. Data Processing Pipeline for Lipid Profiling of Carotid Atherosclerotic Plaque with Mass Spectrometry Imaging. J Am Soc Mass Spectrom 30, 1790–1800, (2019).

62. Oesterle, A., Laufs, U. & Liao, J. K. Pleiotropic Effects of Statins on the Cardiovascular System. Circ Res 120, 229–243, (2017).

63. van der Vusse, G. J. Albumin as fatty acid transporter. Drug Metab Pharmacokinet 24, 300–307, (2009).

64. Su, B. et al. A DMS Shotgun Lipidomics Workflow Application to Facilitate High-Throughput, Comprehensive Lipidomics. J Am Soc Mass Spectrom 32, 2655–2663, (2021).

65. Ghorasaini, M. et al. Congruence and Complementarity of Differential Mobility Spectrometry and NMR Spectroscopy for Plasma Lipidomics. Metabolites 12, (2022).

66. Ledderose, C., Heyn, J., Limbeck, E. & Kreth, S. Selection of reliable reference genes for quantitative real-time PCR in human T cells and neutrophils. BMC Res Notes 4, 427, (2011).

67. Love, M. I., Huber, W. & Anders, S. Moderated estimation of fold change and dispersion for RNA-seq data with DESeq2. Genome Biol 15, 550, (2014).

68. Kassambara, A. & Mundt, F. Extract and Visualize the Results of Multivariate Data Analyses. (2020).

69. Gu, Z., Eils, R. & Schlesner, M. Complex heatmaps reveal patterns and correlations in multidimensional genomic data. Bioinformatics 32, 2847–2849, (2016).

70. Yu, G., Wang, L. G., Han, Y. & He, Q. Y. clusterProfiler: an R package for comparing biological themes among gene clusters. OMICS 16, 284–287, (2012).

71. Wishart, D. S. et al. PathBank: a comprehensive pathway database for model organisms. Nucleic Acids Res 48, D470–D478, (2020).

72. Heinz, S. et al. Simple combinations of lineage-determining transcription factors prime cis-regulatory elements required for macrophage and B cell identities. Mol Cell 38, 576–589, (2010).

73. Sinke, L., Cats, D. & Heijmans, B. T. Omixer: multivariate and reproducible sample randomization to proactively counter batch effects in omics studies. Bioinformatics, (2021).

74. Sinke, L., van Iterson, M., Cats, D., Slieker, R. & Heijmans, B. T. DNAmArray: Streamlined workflow for the quality control, normalization, and analysis of Illumina methylation array data. (2019).

75. van Iterson, M. et al. MethylAid: visual and interactive quality control of large Illumina 450k datasets. Bioinformatics 30, 3435–3437, (2014).

76. Zhou, W., Laird, P. W. & Shen, H. Comprehensive characterization, annotation and innovative use of Infinium DNA methylation BeadChip probes. Nucleic Acids Res 45, e22, (2017).

77. Ritchie, M. E. et al. limma powers differential expression analyses for RNA-sequencing and microarray studies. Nucleic Acids Res 43, e47, (2015).

78. Mandala, W., Harawa, V., Munyenyembe, A., Soko, M. & Longwe, H. Optimization of stimulation and staining conditions for intracellular cytokine staining (ICS) for determination of cytokine-producing T cells and monocytes. Curr Res Immunol 2, 184–193, (2021).

